# Carbon nanotube biocompatibility in plants is determined by their surface chemistry

**DOI:** 10.1101/2021.07.29.454380

**Authors:** Eduardo González-Grandío, Gözde S. Demirer, Christopher T. Jackson, Darwin Yang, Markita P. Landry

**Affiliations:** Department of Chemical and Biomolecular Engineering, University of California, Berkeley, Berkeley, CA, USA; Innovative Genomics Institute (IGI), Berkeley, CA, USA; California Institute for Quantitative Biosciences, QB3, University of California, Berkeley, CA, USA; Chan-Zuckerberg Biohub, San Francisco, CA, USA

**Author notes:** Corresponding author: Markita P. Landry. Department of Plant Biology and Genome Center, University of California, Davis, CA, USA.

**Keywords:** plant biotechnology, nanotechnology, DNA delivery, carbon nanotube, RNA sequencing

## Abstract

Agriculture faces significant global challenges including climate change and an increasing food demand due to a growing population. Addressing these challenges will require the adoption of transformative innovations into biotechnology practice, such as nanotechnology. Recently, nanomaterials have emerged as unmatched tools for their use as biosensors, or as biomolecule delivery vehicles. Despite their increasingly prolific use, plant-nanomaterial interactions remain poorly characterized, drawing into question the breadth of their utility and their broader environmental compatibility. Herein, we characterize *Arabidopsis thaliana* transcriptional response to single walled carbon nanotubes (SWNTs) with two different surface chemistries commonly used for biosensing and nucleic acid delivery: oligonucleotide adsorbed-pristine SWNTs, and polyethyleneimine-SWNTs loaded with plasmid DNA (PEI-SWNTs), both introduced by leaf infiltration. We observed that SWNTs elicit a mild stress response almost undistinguishable from the infiltration process, indicating that these nanomaterials are well-tolerated by the plant. However, PEI-SWNTs induce a much larger transcriptional reprogramming that involves stress, immunity, and senescence responses. PEI-SWNT-induced transcriptional profile is very similar to that of mutant plants displaying a constitutive immune response or treated with stress-priming agrochemicals. We selected molecular markers from our transcriptomic analysis and identified PEI as the main cause of this reaction. We show that PEI-SWNT response is concentration-dependent and, when persistent over time, leads to cell death. We probed a panel of PEI variant-functionalized SWNTs across two plant species and identified biocompatible SWNT surface functionalizations. Our results highlight the importance of nanoparticle surface chemistry on their biocompatibility and will facilitate the use of functionalized nanomaterials for agricultural improvement.

**Significance statement:** Nanomaterials can be used in agriculture as biosensors to monitor plant health, as fertilizers or growth regulators, and as delivery vehicles for genome engineering reagents to improve crops. However, the interactions between nanoparticles and plant cells are not well understood. Here, we characterize the plant transcriptomic response to single-walled carbon nanotubes (SWNTs) commonly used for sensing and nucleic acid delivery. While SWNTs themselves are well tolerated by plants, SWNTs surface-functionalized with positively charged polymers become toxic and produce cell death. We identify molecular markers of this toxic response to create biocompatible SWNT formulations. These results highlight the significance of nanoparticle surface chemistry, perhaps more than the nanoparticles themselves, on downstream interactions of nanoparticles with the environment.

## Introduction

Given the magnitude of environmental changes driven by climate change, and an increasing human population, plant biotechnology and bioengineering have an important role in providing food security, maintaining global biodiversity and sustainability. Technologies that sense plant stress in real-time, improve crop nutritional capacity, render plants resistant to biotic and abiotic stresses, and enable higher crop yield with fewer resources are central to these efforts^1^.

Recently, nanoscience and nanoengineering technologies have been employed in numerous plant biotechnology applications^1,2^. Nanomaterials exhibit unique and tunable size, shape, and physical, mechanical and optical properties. In plant biotechnology, a broad range of nanoparticles have been used including metal nanoparticles, quantum dots, mesoporous silica nanoparticles, clay nanosheets, DNA nanostructures and carbon nanomaterials such as graphene, carbon nanotubes and carbon dots^2^. Within the scope of plant biotechnology, these nanomaterials have been most commonly used as fertilizers, antimicrobials, sensors, imaging agents, and delivery vehicles for small molecules, genes and proteins for plant genetic engineering applications^2–4^.

To accompany the growing use of nanomaterials in plant science, it is essential to understand nanomaterial-plant interactions and the impact of nanomaterials on plant health and their environment. Recent studies have focused on the phenotypical phytotoxicity of metal- and carbon-based nanoparticles on monocot and dicot crop species of interest^5–8^, yet many of these studies show conflicting and often contradictory results. For instance, nano-TiO_2_ (5 nm) has been shown to accelerate germination of spinach seeds^9^, whereas nano-TiO_2_ (20 nm) did not alter the germination rate of wheat grains^10^. Another metal nanoparticle, nano-ZnO (20 nm, 2000 mg/L), has been shown to inhibit root growth in rapeseed, radish, ryegrass, lettuce, corn and cucumber^11^. Cadmium sulfide quantum dots induce oxidative stress and root lignification in soybean^12^. The most studied carbon-based nanomaterials in plant biotechnology are graphene and multi-walled carbon nanotubes (MWNTs). Cabbage, tomato, spinach and lettuce seeds soaked in 2000 mg/L graphene overnight had inhibited growth and reduced biomass^13^. On the other hand, MWNTs enhanced the germination and seedling root elongation when added to the wheat growth medium at 1000 mg/L^6^, and stimulated the growth of roots and stems in legumes^14^. There are several studies discussing the phenotypical phytotoxicity of single-walled carbon nanotubes (SWNTs) with varying results. Studies in *Arabidopsis thaliana* and rice protoplasts show concentration-dependent cytotoxicity of SWNTs, while high SWNT concentrations (250 mg/L) had no observable effects on *Arabidopsis* leaves^15^. Some studies show that the carboxylated SWNTs (COOH-SWNTs) at 50mg/L induce growth and enhanced biomass of tomato plants^16^, whereas poly-3-aminobenzenesulfonicacid functionalized SWNTs at 1750 mg/L inhibited root growth in tomato^17^. Lastly, SWNTs have been shown to reduce oxidative stress and improve cryopreservation of *Agapanthus praecox* embryogenic calli^18^.

More recently, microarray analysis and RNA sequencing (RNA-seq) methods that leverage the advantages of next-generation sequencing have been used to determine nanomaterial impact on plant health^19–21^. Due to its complete gene sequence and annotation, *Arabidopsis* is the preferred plant species for the transcriptomic analyses. For instance, whole-genome microarray analysis of *Arabidopsis* roots exposed to 100 mg/L nano-ZnO revealed up-regulation of genes involved in biotic and abiotic stress responses and down-regulation of genes involved in cell biosynthesis, electron transport, and energy pathways^22^. Additionally, *Arabidopsis* plants that were exposed to 20 mg/L nano-TiO_2_ had significant repression of phosphate-starvation and root-development genes^23^. Lastly, 50 mg/L MWNT treated tomato roots showed substantial upregulation of genes coding for water-channel proteins and hormone pathways^16^, in accordance with the phenotypical data demonstrating enhanced growth in the presence of MWNTs. These studies provide valuable information at the gene expression level and enable a better understanding of the nanomaterial impact on plant health.

Given the recent surge of SWNT usage in plants for sensing ^24–28^ and biomolecule delivery ^29–32^, it is critical to determine the impact of SWNTs on plant health at the molecular and gene expression level. Analogously, it is also important to pinpoint what component(s) of SWNT nanomaterials are responsible for generating differential gene expression patterns, to reconcile conflicting reports. In this study, we performed RNA-seq analysis of *Arabidopsis* leaves 48 hours after exposure to 50 mg/L SWNTs with two different surface chemistries representing the most commonly used SWNT nanomaterials for plant delivery and sensing applications: oligonucleotide-adsorbed pristine SWNTs, and polyethyleneimine (PEI) conjugated functionalized SWNTs (PEI-SWNTs) loaded with plasmid DNA. Our results revealed that SWNTs produced a mild stress response in plants, nearly indistinguishable from a water-infiltration control, that was well tolerated and did not result in permanent damage. However, PEI-functionalized SWNTs at high concentrations produced an adverse response that was irreversible and resulted in cell death, indicating that PEI is the main cause of biotoxicity. We identified gene markers to probe plant responses to different SWNT surface chemistries and discovered new biocompatible SWNT surface chemistry formulations that will facilitate their use in plants.

## Results

### Functionalized PEI-SWNTs generate a very distinct transcriptional response compared to un-functionalized SWNTs

Given the widespread use of PEI-functionalized nanomaterials for delivery of DNA and RNA in plants^30,32–34^, we sought to investigate the reaction of plant tissues to treatment with PEI-SWNTs versus pristine SWNTs. To this end, we infiltrated *Arabidopsis* leaves with pristine single walled carbon nanotubes used in RNA silencing applications (SWNTs)^30^ and polyethyleneimine-functionalized SWNTs used for plasmid DNA delivery (PEI-SWNTs)^32^. We used *Arabidopsis* as it is a well-characterized model plant for which genomic and detailed gene function information is readily available. Aforementioned pristine SWNT and PEI-SWNT nanoparticles were loaded with single stranded RNA targeting Green Fluorescent Protein (GFP) with no target sequence in the *Arabidopsis* genome, and a GFP-expressing plasmid^35^, respectively. For experiments herein, we used ~25-50 fold higher concentrations of SWNTs and PEI-SWNTs compared to standard concentrations used in biomolecule delivery assays to ensure we would observe a robust transcriptional change. Water-infiltrated plant leaves served as a negative control to distinguish between the SWNT-specific response and the response to the infiltration process itself (Figure 1A). We performed RNA-seq with RNA extracted from leaves 48 hours post infiltration to identify changes in the leaf transcriptomic profile in response to the three treatments, compared to non-infiltrated leaves. We validated the RNA-seq data by measuring the expression changes in 12 selected genes using reverse transcription quantitative polymerase chain reaction (RT-qPCR) (Figure S1). Changes in gene expression measured by RNA-seq and RT-qPCR correlated well, confirming the reliability of the RNA-seq data.

**Figure 1.**
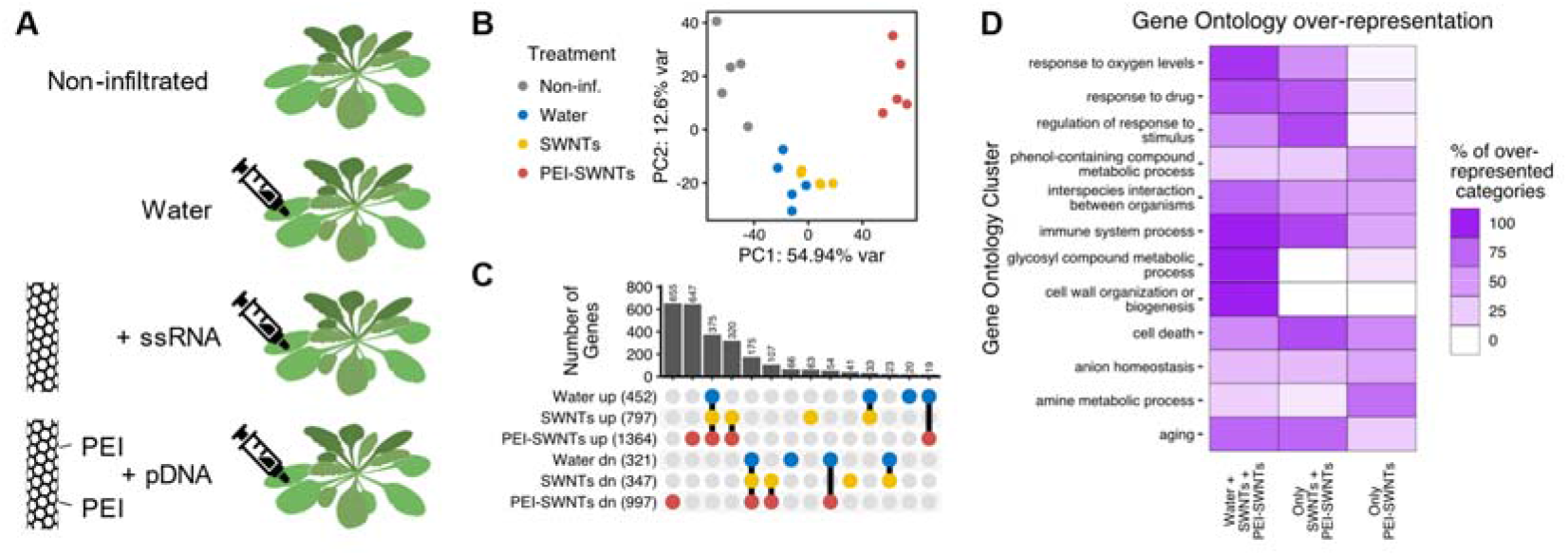
PEI-SWNTs elicit a very distinct transcriptional response compared to that of water or SWNTs. (A) Experimental setup: *Arabidopsis* leaves were infiltrated with ssRNA-adsorbed pristine SWNTs or plasmid-loaded PEI-SWNTs. Non infiltrated and water-infiltrated leaves were used as controls. Samples were collected 48 hours post infiltration. (B) Principal Component Analysis of the transcriptomic profile in response to the different treatments. Each dot represents one biological replicate. (C) Upset plot showing the number of up- or down-regulated genes common or specific to each treatment. Genes with a statistically significant (FDR < 0.05) two-fold change in expression relative to non-infiltrated samples were selected. (D) Functional characterization of genes differentially expressed in the three treatments (375 up, 175 down), specifically in SWNTs and PEI-SWNT samples (320 up, 107 down), and uniquely in PEI-SWNT samples (647 up, 655 down). For a full list of GO terms, see Supplemental Table S2.

RNA-seq data was first analyzed with Principal Component Analysis (PCA) of the whole transcriptomic profile of each sample to assess which treatments induced the largest changes in the plant leaf transcriptome. Surprisingly, PCA analysis revealed that plant response to infiltration with SWNTs is very similar to the response to the water infiltration control, suggesting that SWNTs themselves do not elicit a general transcriptional reprogramming beyond that generated by the infiltration process itself. These results are surprising because they suggest the high concentrations of SWNTs used in our experiments may be biocompatible for use in plants. Conversely, PCA clustering of PEI-SWNTs samples showed that the plant response to PEI-SWNT treatment is distinct relative to SWNT treatment or treatment with water alone (Figure 1B), suggesting that nanoparticle surface chemistry could dictate nanoparticle biocompatibility.

Next, we selected genes that showed a statistically significant (false discovery rate < 0.05) two-fold expression change with respect to the non-infiltrated samples for further analysis. We identified 452, 797, and 1364 up-regulated genes in water, SWNT and PEI-SWNT treated leaf samples, respectively. Conversely, we identified 321, 347, and 997 down-regulated genes in water, SWNT and PEI-SWNT treated leaf samples (Figure 1C, Supplemental Table S1). These results quantitatively confirmed the hypothesis that PEI-SWNTs cause much greater transcriptional reprogramming than the other two treatments. We compared the common genes that respond to the three treatments and observed that 94% (427/452) of the up- and 79% (252/321) of the down-regulated genes in response to water infiltration also changed in SWNT and PEI-SWNT samples, representing genes that change in response to the infiltration process and are independent of a nanomaterial-specific response. Next, to identify biological processes that were present at higher than random fraction when compared to the *Arabidopsis* genome, we performed a Gene Ontology (GO) over-representation analysis. Water/SWNT/PEI-SWNT common genes showed a very significant over-representation of processes related to hypoxia, as expected from injecting a liquid into the leaf tissue that contains large air spaces. Additionally, many GO categories related to cell wall organization, immune response, senescence (aging/programmed cell death), glycosinolate biosynthesis (defense metabolites) and biotic and abiotic stresses were over-represented in this gene set (Figure 1D and Supplemental Table S2A). These results suggest that the infiltration process produces a hypoxia response that triggers other stress-related genes. The fact that we did not observe any phenotypical change in response to water infiltration suggests that this transcriptomic response is insufficient to trigger observable physiological changes in *Arabidopsis* leaves.

We next focused our study on genes that change specifically following treatments with carbon nanotubes, that is, genes that change in leaves infiltrated with SWNT and PEI-SWNT, but do not change following water infiltrations. A majority of genes that change in response to SWNTs also change in response to PEI-SWNTs (76% or 320/422 of up- and 62% or 107/172 of down-regulated genes in SWNTs samples), indicating that these genes are involved in the response to the presence of carbon nanotubes independent of their surface functionalization. GO over-representation analysis showed that genes related to hypoxia, immune response, biotic and abiotic stress responses are also over-represented in this gene set (Figure 1D and Supplemental Table S2B). These results suggest that the presence of carbon nanotubes magnifies the responses to infiltration, because in this analysis we excluded genes that change in all three treatments (e.g., 38 genes in the *cellular* response to *hypoxia* GO category change in all three treatments, while 39 additional genes in the same GO category change only in SWNTs and PEI-SWNTs samples (Supplemental Tables S2A and S2B)). Lastly, certain GO categories including those related to cell death and aging are more over-represented in this gene set when compared to the Water/SWNT/PEI-SWNT genes. These results suggest that the presence of carbon nanotubes exacerbates the plant response triggered by the infiltration process and could indicate that a more advanced response is occurring in these samples at this 48-hour time point. On the other hand, cell wall reorganization and glycosinolate biosynthesis-related processes are not over-represented categories in the SWNT/PEI-SWNT gene set (Figure 1D), indicating that nanotube responses are specific and not a general amplification of the stress caused by infiltration.

PEI-SWNTs induce a much greater transcriptomic reprogramming than water or SWNT treatments, with 647 and 655 up- and down-regulated genes exclusively in PEI-SWNT treated leaves (Figure 1C). In PEI-SWNT treated leaves, we further observed an over-representation of immune system and cell death processes. In plants, a prolonged immune response often leads to programmed cell death^36^, indicating that the plant response to PEI-SWNTs could be more detrimental to treated tissues than less stressful water or SWNT treatments. At the same time, we observe that responses related to aromatic amino acid and secondary metabolite biosynthesis are highly over-represented in PEI-SWNT samples relative to SWNT or water-treated leaves (Figure 1D). Specifically, genes involved in Tryptophan and salicylic acid biosynthesis are specifically over-represented in the PEI-SWNT response. These results indicate that PEI-SWNT specific responses focus on metabolism reconfiguration affecting both primary (i.e., aromatic amino acids) and secondary metabolites involved in defense responses derived from those aromatic amino acids (i.e., salicylic acid)^37^. Taken together, our RNA-seq results suggest that water infiltration alone triggers a mild stress response, one which is highly and at times irreversibly exacerbated by the presence of PEI-SWNTs in a manner not observed following treatment with SWNTs. These results highlight the importance of nanoparticle surface chemistry on nanoparticle biocompatibility in plants.

### PEI-SWNTs strongly up-regulate stress responses and programmed cell death genes, and down-regulate metabolism-related genes

To better characterize the large transcriptional reprogramming of plant leaves treated with PEI-SWNTs, we focused our study on the PEI-SWNT-responding genes that had the highest expression-fold change when compared to water or SWNT treatments, based on the notion that these genes could be the main contributors to the PEI-SWNT specific response. Clustering analyses of gene expression levels in the three treatments revealed a group of 1063 genes that showed very high expression levels in PEI-SWNT treated samples compared to their levels in the other treatments (Figure 2A, Cluster 1). Conversely, 631 genes showed greatly decreased expression levels in PEI-SWNTs when compared to the two other treatments (Figure 2A, Cluster 2). These genes show a very distinct expression profile in PEI-SWNTs compared to the two other treatments, although they can still be up- or down-regulated to a much lesser degree in response to SWNTs or water. We conducted a Gene Set Enrichment Analysis (GSEA)^38^ with these PEI-SWNT specific genes. This powerful analytical method incorporates the degree of gene expression changes in its statistical analysis to produce a quantitative normalized enriched score (NES). This NES represents how a specific gene set (e.g. hypoxia response genes) is enriched in the most up-regulated (positive NES) or most down-regulated (negative NES) genes in a transcriptomic profile used as query (in this case, PEI-SWNT specific genes). GSEA using biological process GOs showed that up-regulated PEI-SWNT-specific genes are enriched in genes related to biotic and abiotic stress responses, defense responses, senescence, and programmed cell death. Down-regulated genes are involved in amino acid and glycosinolate biosynthesis (Figure 2B, Supplemental Table S3A). These results further confirm that PEI-SWNTs induce a programmed cell death response in leaves, in a process that involves metabolism suppression.

**Figure 2.**
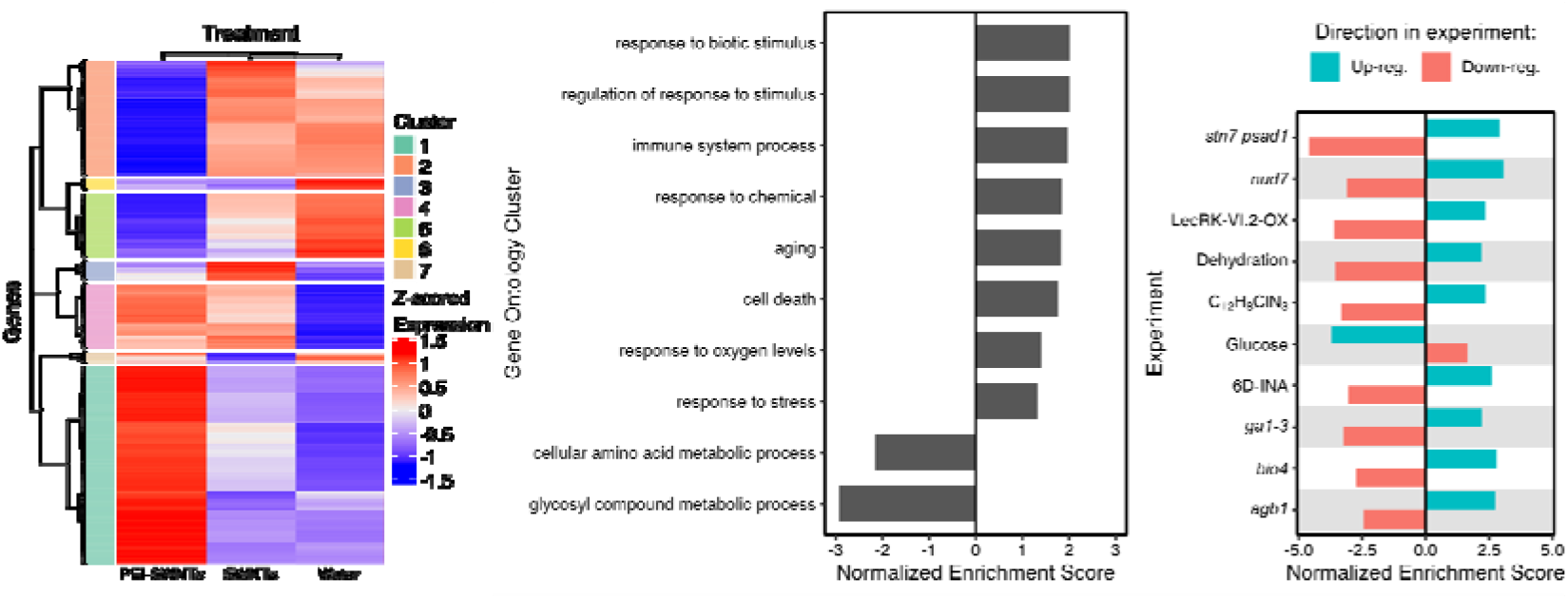
PEI-SWNT responding genes are involved in stress responses, immune system, and programmed cell death. (A) Gene expression heatmap of genes with statistically significant two-fold expression change in at least one of the three treatments, compared to non-infiltrated samples. Clusters 1 and 2 show PEI-SWNT specific up- and down-regulated genes. (B and C) Gene Set Enrichment Analysis of Cluster 1 and 2 genes using Biological Process Gene Ontology categories (B), and Arapath and PlantGSEA databases (C). Up- or down-regulation genes in the original experiment were used as independent gene sets to calculate their normalized enriched score. Details corresponding to each experiment can be found in Supplemental Table S3C.

We also compared the transcriptomic profile of the PEI-SWNT specific genes with other publicly available genome-wide transcriptomic experiments (AraPath and PlantGSEA^39,40^) to find similar profiles that would give us more insight on which processes are involved in the plant response to PEI-SWNTs. This analysis revealed that the expression pattern observed in response to PEI-SWNTs is similar to stress, senescence, and defense responses generated by very different stimuli (Figure 2C, Supplemental Table S3B). Among the most similar transcriptomic profiles, we can find several profiles of *Arabidopsis* mutant plants with altered defense responses, like *stn7 psad1* double mutants defective in photosynthesis acclimation^41^, *nud7* mutants with higher levels of reactive oxygen species^42^, plants over-expressing a lectin receptor kinase that constitutively activates immune responses (LecRK-VI.2)^43^, *bio4* biotin defective mutants that show spontaneous cell death^44^ or *agb1* plant defense signaling heterotrimeric G-protein mutants^45^. Also, PEI-SWNT-induced transcriptomic changes were similar to those of plants treated with chemicals that trigger defense responses like the salicylic acid analog 2,6-dichloroisonicotinic acid^46^ or the slow xenobiotic response inducer 4-chloro-6-methyl-2-phenylpyrimidine^47^. Likewise, the PEI-SWNT specific gene expression pattern was similar to that of plants under dehydration^48^ and opposite to those of plants treated with glucose^49^, suggesting that PEI-SWNT specific genes are involved in responses to metabolic stress (Supplemental Table S3C). Furthermore, GSEA of genes specifically differentially expressed in response to both SWNTs and PEI-SWNTs (Figure 2A, Clusters 4 and 6, up- and down-regulated, respectively) showed highly similar results to the GO over-representation analysis, with response to stress, biotic stimulus and hypoxia being enriched in the up-regulated genes (Supplemental Table 2B). In summary, GSEA using GO and publicly available experiments indicate that the leaf response to treatment with PEI-SWNTs involves stress, immunity, and senescence-related genes.

Finally, we also compared the PEI-SWNT response to profiles of *Agro*-infiltrated *Arabidopsis plants*, the most widely used technique for transient gene expression in plants^50^. We observed a weak but statistically significant enrichment of genes that are up-regulated at 24h and 48h in response to virulent and avirulent *Agrobacterium* strains (Supplemental Table S3D). Up-regulated PEI-SWNT-responding genes that are also up-regulated in response to *Agrobacterium* are mainly involved in detoxification and senescence (e.g., *PCR2*, involved in Zinc detoxification; and *FRK1*, Flagellin/Senescence induced receptor-like kinase1). Conversely, we did not detect an enrichment of agrobacterium down-regulated genes in PEI-SWNT down-regulated genes. Our comparison suggests that the plant response to PEI-SWNT treatment is only partially similar to *Agrobacterium* response.

### PEI is the main cause of toxicity in functionalized PEI-SWNTs

Our results thus far highlight the importance of nanoparticle surface chemistry on inducing differential gene expression patterns in nanoparticle-treated leaves. Specifically, as we observed that PEI-SWNT treatments elicited a much larger transcriptional reprogramming than SWNT treatments, we hypothesized that this difference could be caused by the surface-functionalization of SWNTs with PEI. To better characterize the plant response to PEI-SWNTs, we conducted a more detailed experiment in which we infiltrated increasing concentrations of PEI-SWNTs into *Arabidopsis* leaves. We selected three PEI-SWNT specific genes with large expression changes from the up-regulated Cluster 1 (*PR1, CHX17* and *PAD3*) and the down-regulated Cluster 2 (*AGP41, At3g54830* and *NAI2*), and used these genes as molecular markers to probe the PEI-SWNT concentration-dependent response 48 hours post-infiltration (Figure 3A). Leaves of plants treated with high concentrations of PEI-SWNTs (25 and 50 mg/L) showed some visible damage, especially around the infiltration area, indicating that higher PEI-SWNT concentrations are more toxic to the plant (Figure 3B). We next measured mRNA expression levels of the six marker genes by RT-qPCR in these samples and observed a clear correlation of their expression proportional to PEI-SWNT concentration. Expression of Cluster 1 genes increased with PEI-SWNTs concentration, while expression of Cluster 2 genes decreased (Figure 3C). PEI-SWNTs were loaded with plasmid DNA, but the exogenous DNA does not seem to be the cause of toxicity as leaves infiltrated with no-DNA PEI-SWNTs showed very similar gene expression to plants infiltrated with DNA loaded PEI-SWNTs (Figure 3C). Importantly, COOH-SWNTs, the starting material used for PEI-SWNT functionalization, did not cause any significant change in gene expression even when infiltrated at very high (50 mg/L) concentrations (Figure 3C). These results clearly indicate that PEI is the main causative factor of gene expression changes in plants exposed to PEI-SWNTs.

**Figure 3.**
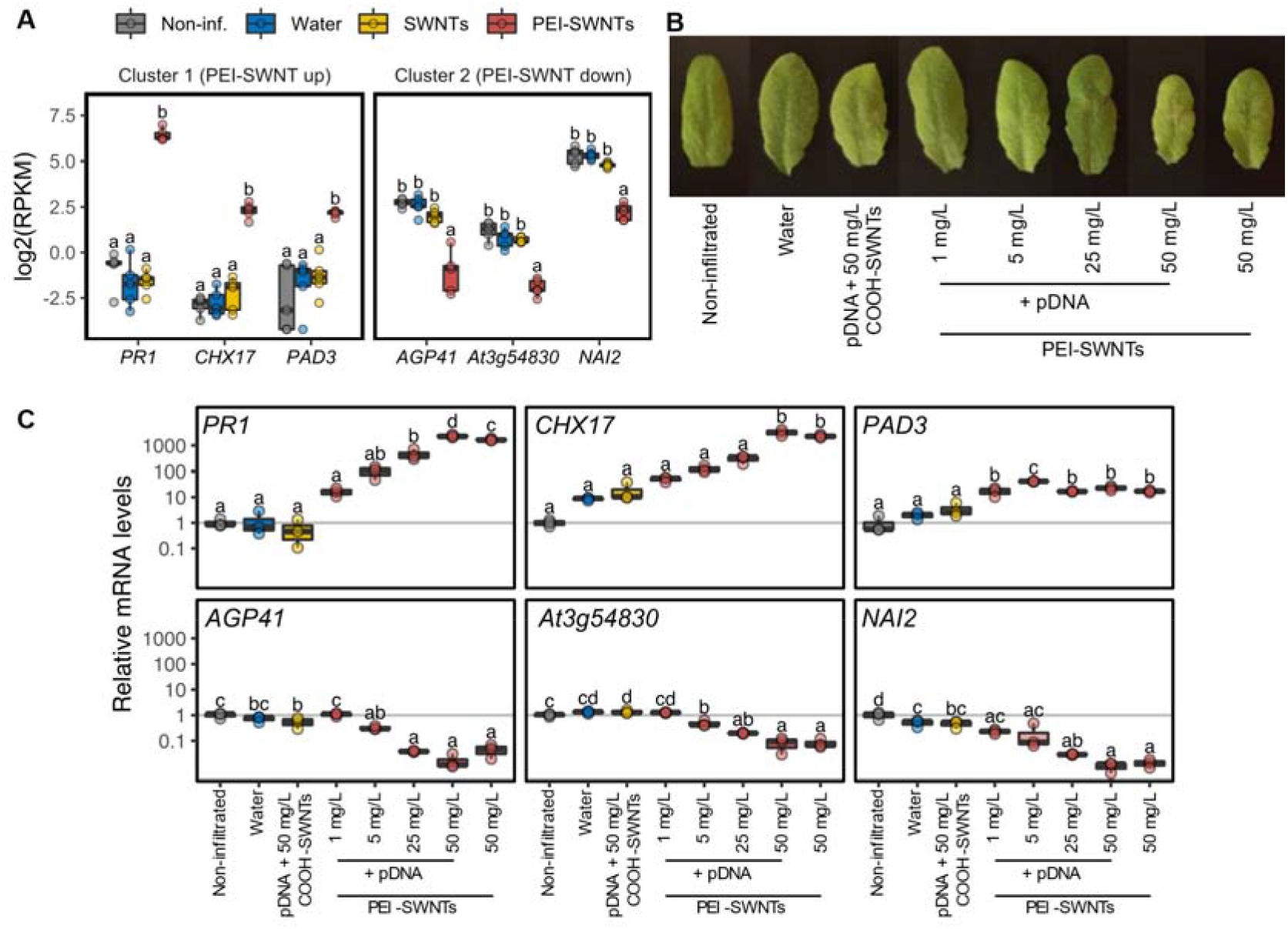
PEI is the main cause of toxicity in PEI-SWNTs. (A) RNA-seq data for three selected genes from Clusters 1 and 2. Each dot represents a biological replicate (n = 5). (B) *Arabidopsis* leaves 48 hours post infiltration with COOH-SWNTs and various concentrations of PEI-SWNTs. (C) mRNA levels of the selected genes measured by RT-qPCR in leaves of plants infiltrated as in (B). The lower and upper hinges of the boxplot correspond to the first and third quartiles, the upper and lower whiskers correspond to the largest value no further than 1.5 times the inter-quartile range. Statistical significance was determined by one-way ANOVA with post-hoc Tukey HSD test. Letters denote significant differences among means (n = 3).

### Persistent PEI-SWNT specific response leads to leaf damage

In our initial experiments, we studied the response to PEI-SWNTs at relatively short times after infiltration (48 hours). To gain better insight into the long-term effects of PEI-SWNTs infiltration, we performed a time course experiment where we infiltrated low (1 mg/L, equivalent to the dose used in standard delivery experiments) and high (50 mg/L) concentrations of PEI-SWNTs, and collected samples 2-, 4-, 6- and 8-days post infiltration (dpi). Again, we detected some leaf damage in 50 mg/L infiltrated samples by the 2-dpi time point. This damage increased as time advanced, with slight leaf chlorosis and evident cell death around the infiltration area at the later time points, indicative of a strong stress response (Figure 4A). These symptoms were not apparent in 1 mg/L PEI-SWNT infiltrated leaves, suggesting that low PEI concentrations do not elicit this toxic response over time. We measured expression levels of the 6 marker genes by RT-qPCR at each time-point. The measured expression patterns were consistent with our previous results at 2-dpi, whereby 50 mg/L PEI-SWNTs induced larger expression changes than 1 mg/L PEI-SWNTs. Expression of up-regulated genes (*PR1, CHX17* and *PAD3*) decreased over time for both treatments, with 50 mg/L samples always showing higher levels than 1 mg/L. At 6-dpi, expression levels of these genes in 1 mg/L samples returned to values close to basal levels (non-infiltrated samples) indicating that the response to low concentration of PEI-SWNTs had subsided. Conversely, even at 8-dpi, expression in 50 mg/L samples was still higher than non-infiltrated samples. RT-qPCR analysis of down-regulated genes (*AGP41, At3g54830* and *NAI2*) showed similar trends, albeit with a more variable expression pattern at later timepoints. *AGP41* showed a stable pattern, with greater downregulation in 50 mg/L samples than in 1 mg/L samples at every time point, recovering to basal levels in 1mg/L samples at 8-dpi. Meanwhile, *At3g54830* and *NAI2* were down-regulated at 2-dpi, up-regulated at 4-dpi in 50 mg/L samples, and down-regulated again at later time points. Their expression in 1 mg/L samples decreased at 4-dpi and recovered to basal levels at 6- and 8-dpi (Figure 4B). These results suggest that a prolonged activation of PEI-SWNT specific genes generates a more severe and irreversible programmed cell death response.

**Figure 4.**
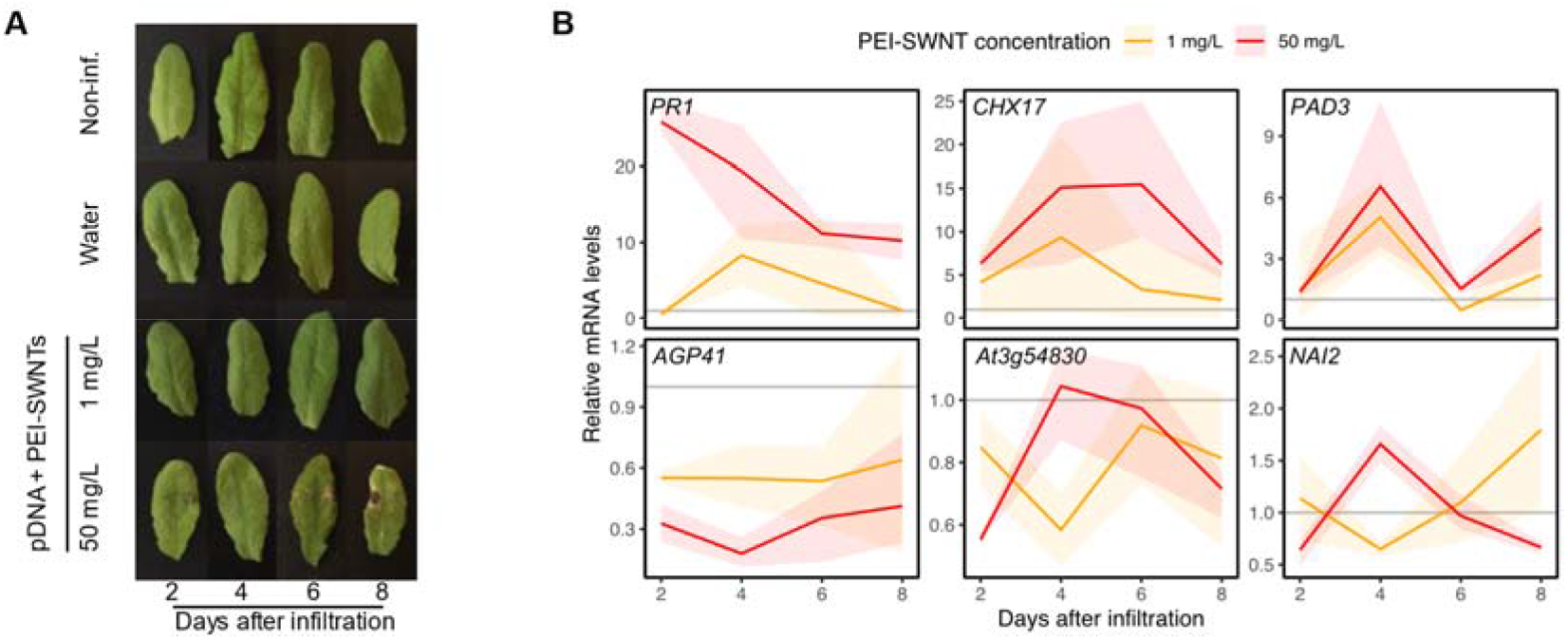
Persistent PEI-SWNT specific gene response leads to severe leaf damage. (A) *Arabidopsis* leaves infiltrated with different PEI-SWNTs concentrations. Images taken two, four, six and eight dpi. (B) mRNA levels of selected marker genes at different time points after infiltration. mRNA levels are normalized to non-infiltrated samples at the corresponding time point (represented by a grey line). Each colored line represents the average mRNA levels at each time point and the faded band represents a 95% confidence interval (n =3).

### Identification of biocompatible functionalized SWNTs

Our results demonstrate that toxicity generated by PEI in PEI-SWNTs can become a limiting factor when high concentrations of PEI-SWNTs are required to ensure efficient biomolecule delivery. To find more biocompatible SWNT surface chemistries we infiltrated *Arabidopsis* leaves with several PEI polymer variants covalently bound to SWNTs with similar zeta potential and DNA binding capabilities to PEI -SWNTs^51^. We infiltrated 50mg/L of low molecular weight linear PEI (800 Da; L-PEI 800), hydrophobically modified branched PEI (25-30 kDa; H-PEI) and high molecular weight branched PEI (750 kDa; PEI 750k). We included unfunctionalized COOH-SWNTs and branched PEI-SWNTs (25 kDa; PEI) as negative and positive toxicity controls, respectively (Figure 5A). We measured mRNA levels of the 6 toxicity marker genes and found that L-PEI 800 showed an expression pattern very similar to COOH-SWNTs (Figure 5B), indicating that SWNTs functionalized with this polymer are more biocompatible than PEI-SWNTs. H-PEI showed a similar pattern to PEI, while PEI 750k showed a lower degree of toxicity.

**Figure 5.**
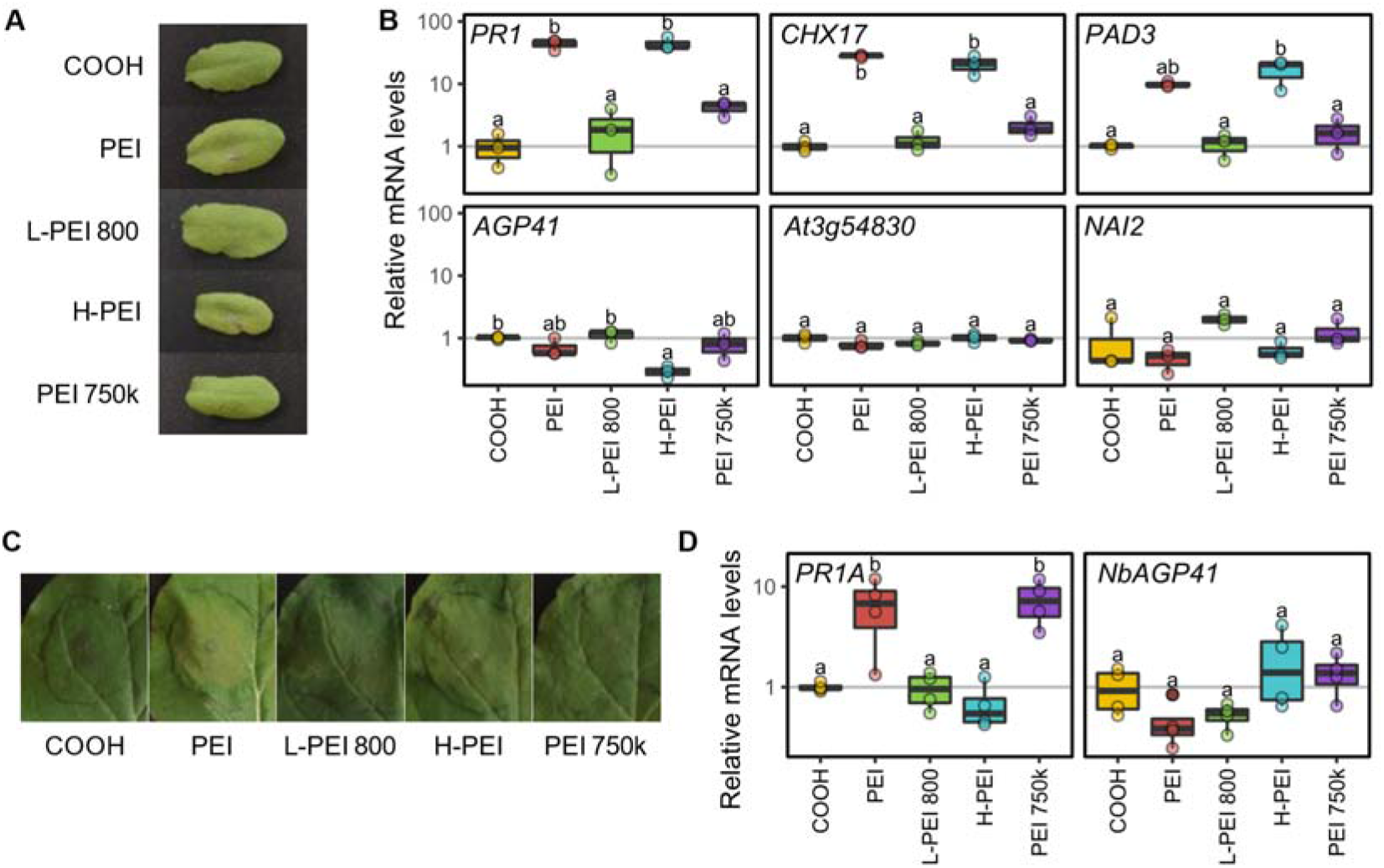
Low molecular weight linear PEI functionalized SWNTs are more biocompatible than PEI-SWNTs. (A) *Arabidopsis* leaves infiltrated with COOH-SWNTs and SWNTs functionalized with different polymers. (B) mRNA levels of the selected marker genes measured by RT-qPCR in leaves of plants infiltrated as in (A). (C) *Nicotiana benthamiana* leaves infiltrated with the same SWNT preparation as in (A). (D) mRNA levels of *Nicotiana PR1* and *AGP41* ortholog genes measured by RT-qPCR. Leaf images taken two days after infiltration. mRNA levels are normalized to COOH-SWNTs. The lower and upper hinges of the boxplot correspond to the first and third quartiles, the upper and lower whiskers correspond to the largest value no further than 1.5 times the inter-quartile range. Statistical significance was determined by a one-way ANOVA with post-hoc Tukey HSD test. Letters denote significant differences among means (n = 3 for *Arabidopsis* and n = 4 for *Nicotiana*).

Lastly, to probe whether our biocompatibility results from *Arabidopsis* would be applicable to other plant species, we tested the response of *Nicotiana benthamiana* to our polymer-SWNTs. Nicotiana is another model plant species that was used to develop the PEI-SWNT delivery technique^32^, and is an extremophile^52^ that tolerates abiotic stresses better than *Arabidopsis*. We identified the Nicotiana orthologs of *Arabidopsis PR1* and *AGP41* (*PR1A* and *NbAGP41*) and used them as molecular markers of toxicity. With the same panel of polymer-SWNTs, we measured their mRNA levels 2-dpi. We only observed slight chlorosis in PEI-SWNT infiltrated samples (Figure 5C). As in *Arabidopsis*, L-PEI 800-SWNTs showed an expression pattern very similar to that of COOH-SWNTs, indicating that this polymer also does not elicit a further stress response in *Nicotiana* (Figure 5D). Surprisingly, *PR1A* levels after H-PEI- and PEI 750k-SWNTs infiltration were the opposite of what we observed in *Arabidopsis*: H-PEI-SWNTs seemed to be well tolerated in *Nicotiana*, while PEI 750k-SWNTs induced a similar response to PEI-SWNTs. These results suggest that there might exist species-specific responses to different SWNT formulations, highlighting the importance of adapting SWNTs functionalization to the species being targeted to maximize biocompatibility. We propose that the data obtained in our RNA-seq experiment can be used as a starting point for the identification of molecular markers that guide generation of nanoparticle-based biotechnologies with enhanced biocompatibility in plants.

## Discussion

Biomolecule delivery and metabolite sensing by carbon-based nanoparticles, such as SWNTs, has emerged as a promising technology to monitor plant health^26,28^, enable precise gene downregulation (siRNA delivery^30,33^) and transient gene expression (plasmid DNA delivery by PEI-functionalized nanomaterials^32,34^). One main advantage of nanoparticle-based biomolecule delivery is that the mechanism by which these nanomaterials penetrate the cells is species-independent^53^, and thus, they can be used in plant species recalcitrant to current delivery methods. Present knowledge on plant-nanomaterial interactions focuses mainly on nanomaterials applied to soil or plant growth medium^1,54^. Understanding the mechanistic basis of how plants react to nanomaterials such as PEI-SWNTs once they are inside the cell and identifying the key components of nanoparticles that generate bioincompatible outcomes is crucial to inform rational improvement of these new technologies.

In this work we implement high-throughput sequencing to provide evidence that the infiltration of SWNTs produces a stress response in plants related to hypoxia, immune system activation, and senescence. This reaction is very similar to the response to the infiltration process itself and is well-tolerated by the endogenous detoxifying mechanisms of the plant. Previous studies in which much higher SWNT concentrations were infiltrated into *Arabidopsis* leaves (250 mg/L, five-fold the highest concentration used in our study) did not detect any macroscopic change either, suggesting that SWNTs can be well tolerated by plants^15^. Even at these high concentrations, transient activation of reactive oxygen scavenging mechanisms seem to be sufficient to eliminate any temporal toxic effects caused by nanomaterials^15^.

However, these stress responses are greatly exacerbated by the presence of PEI in functionalized SWNTs, the main cause of toxicity when high concentrations of PEI-SWNTs are infiltrated. To our knowledge, no PEI toxicity studies have been reported in plants. We observed that PEI-SWNT specific responses are concentration-dependent and when sustained over time, lead to visible tissue damage in tissues exposed to a high concentration of PEI-SWNTs. PEI exerts a wider transcriptional reprogramming that leads to metabolism suppression and programmed cell death in the infiltrated areas. These responses are only partially similar to those elicited by other commonly used nucleic acid delivery techniques, such as agroinfiltration^50^, indicating that these are specific to nanomaterial functionalization. Most importantly, these results highlight that nanoparticle surface chemistry, moreso than the nanoparticle itself, can drive the biocompatibility or lack thereof of SWNT-based plant biotechnologies, a finding that may be extendible to other nanoparticle types.

Generally, nanoparticle-mediated biomolecule delivery is less efficient than conventional biotic delivery methods such as *Agro*-infiltration. Our results highlight that this lower efficiency cannot be overcome by simply increasing the amount of delivered PEI functionalized nanoparticles, as they show toxic effects when plants are treated with high concentrations. Thus, to identify more biocompatible SWNT preparations, we measured the response of marker genes identified in our transcriptomic analysis to SWNTs functionalized with a panel of cationic polymers. We observed that in *Arabidopsis*, low molecular weight linear PEI resulted in a lower toxicity response, suggesting this functionalization as a viable alternative to the currently used PEI. It is interesting that the low stress response to L-PEI 800 observed in *Arabidopsis* is conserved in *Nicotiana*, suggesting that this polymer’s uniquely small size and lower amine density plays a key role in not triggering stress response pathways across plant species. Similarly, the response to PEI-750k is conserved across both species tested. Counterintuitively, we attribute the relatively low toxicity of this polymer to its large size, which may limit its ability to internalize in cells. Separately, the species-specific response to H-PEI supports the importance of hydrophobicity in nanomaterial interactions with aqueous cellular components. We hypothesize that the lack of response in *Nicotiana* could be attributed to alternate mechanisms for managing hydrophobic substances or a response through a different pathway that does not involve the *PR1A* gene. For instance, proteins and other biomolecules can adsorb to nanomaterials forming a bio-corona^56–58^, and its composition can change depending on the nanomaterial^59^ and its functionalization^60^. Currently, studies of nanomaterial-corona formation in plants are scarce^61–63^. Species-specific bio-corona formation could account for the difference in stress response we observed, and further highlights the need for more studies using different nanomaterials and plant species to inform tailored nanomaterial functionalization.

Interestingly, when these stress responses were activated at lower levels by low concentrations of PEI-SWNTs, they were well tolerated by plants after several days. It remains to be studied if this temporal response could prime plants to better resist later biotic or abiotic stresses. Indeed, the observed PEI-SWNT response is similar to the one elicited by defense-priming agrichemicals used to confer long-lasting resistance to biotic^46^ and abiotic stresses^47^. This opens the possibility of using low PEI-SWNT concentrations as stress-priming treatments. In fact, silica nanoparticles have recently been shown to enhance disease resistance through salicylic acid-mediated systemic acquired resistance^64^. Further work studying these and longer-term effects of PEI-SWNTs on plants are needed to explore this possibility.

The framework described in this work (transcriptomic profiling, marker identification and concentration/time/functionalization-dependent response validation) could be adapted to study the plant responses to other agronomically-relevant nanomaterials with promising applications in different plant species. Once nanomaterial-plant interactions are better characterized, rational design of more biocompatible functionalized nanomaterials can be achieved for a broad range of plant biotechnology applications.

## Materials & Methods

### Nanomaterial preparation

ssRNA-adsorbed SWNTs were prepared as described in^30^. Plasmid DNA adsorbed PEI-SWNTs were prepared as described in^29^. Other polymer-SWNTs were prepared as described in^51^.

### RNA-seq sample collection and preparation

Wild-type Columbia-0 *Arabidopsis thaliana* and wild-type *Nicotiana benthamiana* plants were grown in the greenhouse and in a HiPoint 740FHLED growth chamber, respectively, under the following conditions: 24°C high and 21°C low, 16 h light and 8 h dark, and 70% average humidity. Young leaves^65^ of 6-week old wild-type Col-0 *Arabidopsis* plants in vegetative stage were selected to be fully infiltrated (approx. vol. 40 μL) with a 1 mL needleless syringe (BD, cat. no. 14-823-434) loaded with water, COOH-SWNTs (50 mg/L), SWNTs (50 mg/L), PEI-SWNTs (50 mg/L) or the panel of polymer-SWNTs (50 mg/L). The third and fourth leaves of 4-week-old *Nicotiana* plants were infiltrated in the same way. Two days after infiltration, five biological replicates containing four *Arabidopsis* leaves of each treatment from different plants and from non-treated plants were collected. For *Nicotiana*, a 1 cm by 1 cm infiltrated area was collected for each biological replicate. Samples were collected in a 2 mL Eppendorf tube with two 3.2 mm chrome steel beads (RPI, cat. no. 9840) and flash frozen in liquid N_2_ immediately. Frozen samples were ground in a Mini-beadbeater (Biospec Products, cat. no. 3110Bx) tissue homogenizer for 5 seconds at 25 Hz frequency, twice.

RNA was extracted using RNeasy Plant Mini Kit (Qiagen cat. 74904) using RNase-Free DNase (Qiagen cat. 79254) following manufacturer instructions. Total RNA concentration was measured using the Qubit™ RNA BR Assay Kit (Thermo Fisher). RNA quality was checked using a 2100 Bioanalyzer with RNA 6000 Nano Kit (Agilent). RNA integrity number (RIN) scores were confirmed to be >8. Libraries were prepared using Kapa Biosystems library preparation kit with mRNA selection with poly-A magnetic beads. Libraries were pooled and sequenced on an Illumina NovaSeq S4 flow cell in a NovaSeq 6000 Platform with 150 paired end reads. On average, 29.5 million reads per sample were obtained. Sequencing data discussed in this publication have been deposited in NCBI’s Gene Expression Omnibus^66^ and are accessible through GEO Series accession number GSE172278.

### Sequencing data analysis

Raw reads were pre-processed using FastQC, and Trimmomatic was used to trim low quality reads^67^. HI-SAT2 was used to map the reads to the *Arabidopsis* genome (TAIR10) using default values^68^. FeatureCounts was used to assign reads to *Arabidopsis* transcripts (TAIR10)^69^. These steps were performed using the public server at usegalaxy.org^70^. Further analysis was performed in R^71^. edgeR was used to identify differentially expressed genes^72^. Briefly, genes with average RPKM values below one were removed from the analysis, then a model matrix was built to compare all the treatments against the non-infiltrated samples, and quasi-likelihood dispersion estimation and hypothesis testing were performed. Genes with a statistically significant (FDR < 0.05) two-fold expression change in at least one of the treatments with respect to the untreated samples were selected for further analysis. The ggupset R package was used for Figure 1B. Gene Ontology Enrichment Analyses were performed using the clusterProfiler package^73^ with GO annotations from org.At.tair.db version 3.11.4. GSEA analysis was performed using the fgsea package^74^ with GO annotations or experimental databases Arapath^39^ and PlantGSEA^40^. GO were aggregated by hierarchical clustering using relevance semantic similarity^75^ and the most common ancestor term with the highest information content was selected as the representative GO term for each cluster. The 10 top experiments in Arapath and PlantGSEA databases with the highest positive NES in PEI-SWNT up-regulated genes and negative NES in PEI-SWNT down-regulated genes were selected for analysis. To identify gene clusters, row Z-scored log2(fold change) values were clustered using Pearson’s correlation as distance and a complete agglomeration method, and represented using ComplexHeatmap package^76^.

### RNA expression analyses

RNA was extracted from 100 mg of ground leaves following the protocol described in^77^ with certain modifications. We used TRIzol (Thermo cat. 15596026) instead of phenol and Phasemaker tubes were used to separate the aqueous phase (Thermo cat. A33248). 10 μg of total RNA were treated with TURBO DNase I (Thermo cat. AM2238) and cDNA was synthetized using 1 μg RNA using the High-Capacity cDNA Reverse Transcription Kit (Thermo cat. 4368814). PowerUP SYBR Green Master Mix (Thermo cat. A25741) was used for RT-qPCR using three technical replicates per reaction in a CFX96 Touch Real-Time PCR Detection System (Biorad). Primers are described in Table S4. *SAND* and *EF1a* genes were used as reference for *Arabidopsis* and *Nicotiana*, respectively^78^. For experiments in *Nicotiana* we identified the closest orthologs of *Arabidopsis* genes using their protein sequence as input in the solgenomics BLAST tool against the *Nicotiana* benthamiana genome^79^. We used the sequence of Niben101Scf00107g03008.1 (*PR1A*) and Niben101scf01817g00015.1 (*NbAGP41*).

---

**Supplemental Figure S1.**
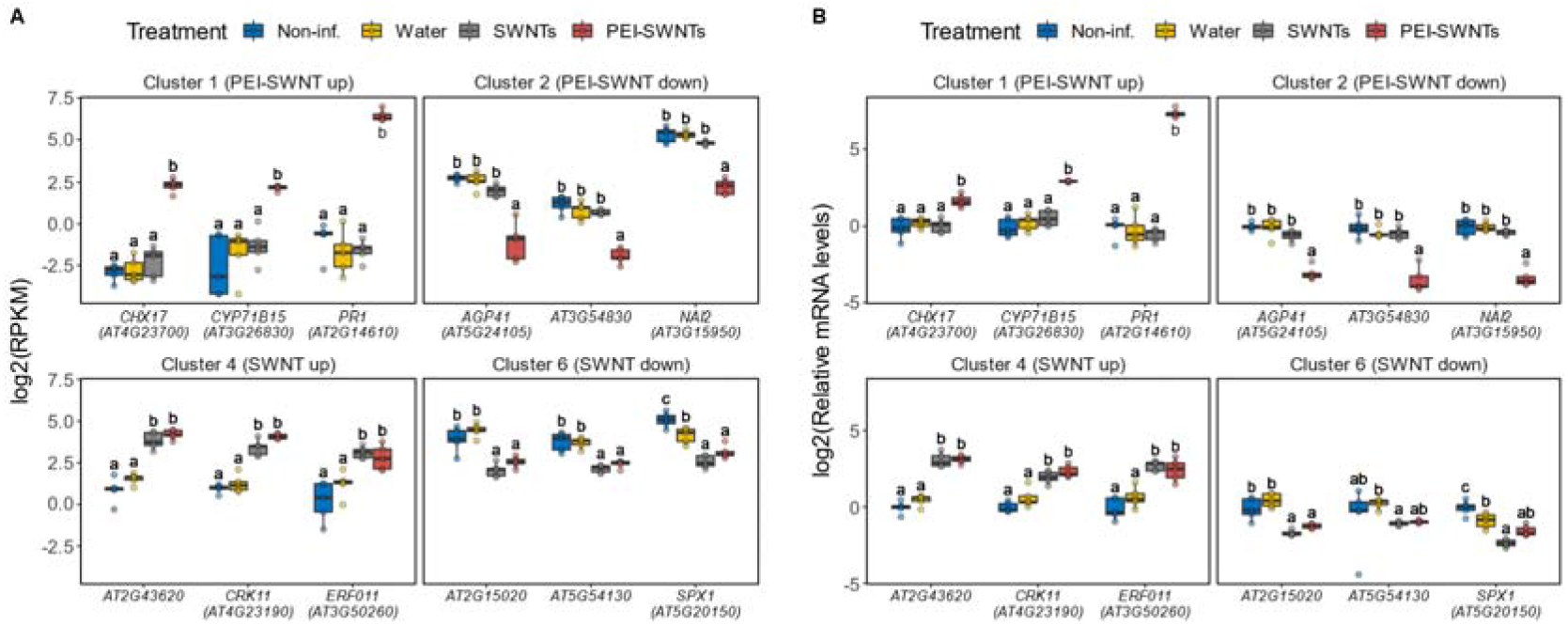
Validation of RNA-seq data by RT-qPCR. (A) Normalized log2(RPKM) values for selected genes from Clusters 1 and 2 (PEI-SWNT specific genes) and Clusters 4 and 6 (SWNT specific genes) from RNA-seq data. (B) mRNA levels of same genes as in (A) measured by RT-qPCR. The lower and upper hinges of the boxplot correspond to the first and third quartiles, the upper and lower whiskers correspond to the largest value no further than 1.5 times the inter-quartile range. Statistical significance was determined by a one-way ANOVA with post-hoc Tukey HSD test. Letters denote significant differences among means (n = 5 in A and n = 3 in B).

## Acknowledgements

We thank Frederic Bouche, parkjisun from Noun Project, and Natalie Goh for the graphics used in Figure 1A. The GFP plasmid was obtained from the Sheen Lab (Harvard Medical School). We thank BASF for providing the L-PEI-800 and H-PEI polymers. We acknowledge support a Burroughs Wellcome Fund Career Award at the Scientific Interface (CASI) (to M.P.L.), a Dreyfus foundation award (to M.P.L.), a Beckman Foundation Young Investigator Award (to M.P.L.), an NIH MIRA award (to M.P.L.), an NSF CAREER award (to M.P.L), an NSF CBET award (to M.P.L.), an NSF CGEM award (to M.P.L.), a FFAR Young Investigator award (to M.P.L.), a CZI investigator award (to M.P.L), a Sloan Foundation Award (to M.P.L.), a USDA BBT EAGER award (to M.P.L), a USDA NIFA Award (to M.P.L), a Moore Foundation Award (to M.P.L.), and a DOE office of Science grant with award number DE-SC0020366 (to M.P.L.) and NSF GRFP (to C.T.J.). M.P.L. is a Chan Zuckerberg Biohub investigator, a Hellen Wills Neuroscience Institute Investigator, and an IGI Investigator.

